# BRACE: A novel Bayesian-based imputation approach for dimension reduction analysis of alternative splicing at single-cell resolution

**DOI:** 10.1101/2024.08.01.606201

**Authors:** Sean Wen

## Abstract

Bayesian approach is a powerful tool to solve challenging questions in life sciences. One such area of life sciences in which Bayesian approach has seen an increased utility in the recent years is single-cell biology. Alternative splicing represents an additional layer of complexity underlying gene expression profile that has the potential to reveal insights into the biological mechanisms underpinning heath and disease states. Dimension reduction analysis is the cornerstone of RNA-sequencing analysis and has the ability to guide selection of candidate biomarkers based on segregation of sample groups. Nevertheless, dimension reduction analysis at single-cell resolution remains a significant challenge for alternative splicing datasets, and therefore hitherto preclude the assessment of candidate isoforms. Here, we introduce BRACE (a Bayesian-based imputation approach for dimension Reduction Analysis of alternative splicing at single-CEll resolution). We demonstrated our Bayesian approach represents an improvement over existing methods for imputing missing percent spliced-in values, and subsequently applied our approach for the dimension reduction analysis of alternative splicing events at single-cell resolution. We further demonstrated the application of our Bayesian approach over a range of single-cell datasets with increasing complexity, namely cell populations that are transcriptomically distinct, similar, and heterogenous. We anticipate our approach to enable assessment and selection of cell state- or disease-specific biomarkers for downstream experimental validation.

## 1 Introduction

Alternative splicing is the process of generating different isoforms from the same gene by utilising different combinations of exons and introns. This process generates isoform diversity, and by extension, protein and phenotype diversity (Matlin, Clark, & Smith, 2005). Alternative splicing plays a role in both health and disease states. In particular, aberrant splicing mechanism is prevalent in diseases such as cancer whereby alternative splicing is dysregulated by mutations on canonical splice sites, mutations that generate cryptic splice sites, mutations affecting genes encoding for splicing factors, and dysregulated expression of RNA-binding proteins (Alsafadi et al., 2016; Kahles et al., 2018; E. Wang et al., 2019).

While alternative splicing analysis is an emerging area of research with potential therapeutic implications, both bulk and single-cell RNA-sequencing analyses have hitherto focused on gene expression profiling (Roy et al., 2021; G. Wang et al., 2022; Wen & Leong, 2019). This is not surprising given that gene expression analysis framework, in particular single-cell analysis, is more established compared to alternative splicing (Satija, Farrell, Gennert, Schier, & Regev, 2015). The single-cell gene expression analysis framework typically consists of gene expression quantification, dimension reduction analysis, cell type assignment, differential expression analysis, and pathway enrichment analysis.

Dimension reduction analysis is helpful for several purpose. Unsupervised clustering based on highly variable genes enables identification of cell clusters (populations) and subsequent assignment of cell type identity based on cell type-specific gene expression signatures (G. Wang et al., 2022). On the other hand, supervised clustering assesses the ability of candidate genes, typically identified from differential gene expression analysis, to segregate different cell states, such as between health and diseased states. This aids identification of candidate genes for downstream experimental validation (Wen & Leong, 2019).

Recently, several single-cell alternative splicing frameworks have been developed that includes splicing detection, quantification, dimension reduction analysis, and differential splicing analysis (Song et al., 2017; Wen, Mead, & Thongjuea, 2020, 2023). Nevertheless, dimension reduction analysis of single-cell alternative splicing datasets faced significant challenges compared to single-cell gene expression datasets. Notably, the challenge of missing values or dropouts in single-cell alternative splicing dataset hampers robust dimension reduction analysis.

In single-cell gene expression datasets, the expression values range from 0 to theoretically high values. Here, a 0 value may represent a technical dropout whereby the mRNA was not sufficiently captured during library preparation and therefore not represented in the sequencing library. The 0 value may also simply indicate that the cell does not express the gene. Nevertheless, regardless of the reason behind the 0 values, dimension reduction analysis may still be proceeded as these technical dropouts or non-expressed genes in the expression matrix are coded as 0.

On the other hand, in single-cell alternative splicing datasets, the values are represented in percent spliced-in (PSI), and the values range from 0 to 100. In contrast to gene expression, a PSI value of 0 does not represent technical or biological dropout. Instead, a PSI value of 0 indicates that the alternative exon is spliced out whereas a value of 100 indicates that the alternative exon is spliced in. PSI values between 0 and 100 indicate varying degree of alternative exon inclusion. In any case, in order for the PSI values to be computed, the isoform, and by extension gene, would need to be expressed. Therefore, isoforms that are not expressed are represented by missing values (“NA”) in the PSI matrix. These missing values preclude downstream dimension reduction analysis.

One possible way to perform dimension reduction analysis on single-cell alternative splicing datasets is to impute these missing values. Current method for imputation of these missing values is a random approach (Song et al., 2017; Wen et al., 2023). Specifically, missing values are randomly imputed with any values between 0 and 100. While published studies have demonstrated successful segregation of different cell populations using this approach, these studies only investigated cell populations that are transcriptomically distinct from one another. This approach has yet to be demonstrated on transcriptomically more complex datasets such as cell populations that are transcriptomically similar or heterogenous.

Therefore, in this study, we developed a novel Bayesian-based imputation method for PSI estimation and demonstrated its application on dimension reduction analysis of single-cell alternative splicing dataset to enable dimension reduction analysis across a range of datasets with differing complexity. To this end, we benchmarked our Bayesian imputation approach against the random imputation approach, and demonstrated the application of both approaches on cell populations of increasing complexity, namely cell populations that were transcriptomically distinct, similar, and heterogenous. It is noteworthy that our approach is applicable to isoforms quantified at the exon-level from short-read RNA-sequencing data (henceforth “splicing event”) generated from plate-based library preparation methods, such as Smart-seq2 and SMARTer protocols (Picelli et al., 2013; Verboom et al., 2019).

We anticipate our Bayesian imputation approach to be helpful in assessing the ability of differentially spliced exons to distinguish different cell populations, and consequently guide the selection of cell state- or disease-specific biomarkers for downstream experimental validation.

## 2 Method

### 2.1 Percent spliced-in (PSI) quantification

Alternative splicing was quantified for seven exon-level splicing event types, namely skipped-exon (SE), mutually exclusive exons (MXE), retained intron (RI), alternative 5’ and 3’ splice sites (A5SS, A3SS), and alternative first and last exons (AFE, ALE) as previously described (Wen et al., 2023). We further improved the detection of splicing events prior to PSI quantification. Firstly, we only included cryptic A5SS and A3SS defined as alternative splice sites located within 100bp of the canonical splice sites. Secondly, we only included *bona fide* AFE and ALE defined as alternative splice sites located beyond 100bp of the canonical splice sites. This will preclude cryptic A5SS and A3SS from being misclassified as AFE and ALE respectively. Cryptic A5SS and A3SS and *bona fide* AFE and ALE may be detected using the *SubsetCrypticSS* and *RemoveCrypticSS* function, respectively, of the *MARVEL* R package.

The unit of splicing measurement is percent spliced-in (PSI), and it represents the percentage of splice junction reads supporting the inclusion (splicing in) of the alternative exon divided by the total number of reads supporting the inclusion or exclusion (splicing out) of the alternative exon as represented in equations 1-3 as follows:

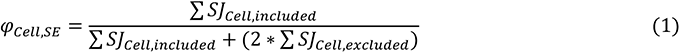

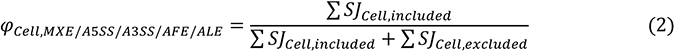

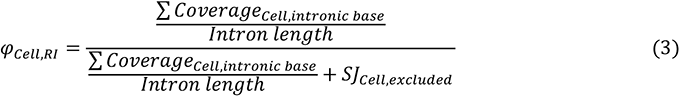

Only splicing events supported by at least 10 splice junction reads were eligible for PSI computation. PSI computation may be performed using the *ComputePSI* function.

### 2.2 Bayesian imputation

One common approach for imputation prior to dimension reduction analysis is replacing the missing values of a given variable with its mean across all samples. This is synonymous with replacing missing PSI values of a given splicing event with its mean across all cells. The main drawback of mean imputation is that it leads to shrinkage of standard deviation or variance, and as a consequence, underestimates the heterogeneity of the splicing patterns across the cell population. It is noteworthy that even a phenotypically homogenous cell population may consist of sub-populations with distinct splicing patterns (Shalek et al., 2013). Moreover, mean imputation does not preserve the relationship between the different splicing events, such as splicing events correlated to one another. Notably, functionally related genes, and by extension, their isoforms are often co-regulated (Trapnell et al., 2014). Nevertheless, mean imputation is still useful when the number of samples with missing values are negligible. However, most splicing events are only expressed in a small percentage of cells, thus exacerbating the drawbacks of mean imputation.

To circumvent the limitations of mean imputation, we proposed a Bayesian method of imputation of missing values pervasive in single-cell alternative splicing datasets. A Bayesian framework consists of three components, namely the likelihood, prior, and posterior probability. Here, we defined the single-cell PSI values as the likelihood and population-level PSI values as the prior.

Consider an alternative 3’ splice site (A3SS) splicing event in a single cell as an example. The likelihood component is the observed PSI value for the single cell as follows in equation (1). The numerator is total number of splice junctions (SJ) supporting the inclusion (splicing in) of the alternative exon, i.e. isoform molecules that express the alternative exon. The denominator is the total number of splice junctions supporting the inclusion (splicing in) or exclusion (splicing out) of the alternative exon, i.e, total coverage at that site across all isoform molecules.

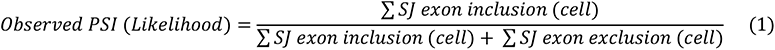

The prior consists of pseudocounts. First, the total number of splice junctions supporting the inclusion of the alternative exon across the entire cell population is computed as shown in mathematical notation (2). Next, the total number of splice junctions supporting the inclusion or the exclusion of the alternative exon across the entire cell population is computed as shown in mathematical notation (3).

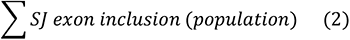

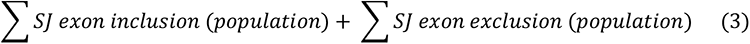

The pseudocount (prior) to be added to the numerator of the likelihood function in equation (1) is the value resulting from the division of mathematical notation (2) by mathematical notation (3) as shown in equation (4) below. This pseudocount value represents an approximation of the population average number of splicing junctions supporting the inclusion of the alternative exon.

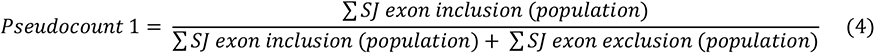

On the other hand, the pseudocount (prior) to be added to the denominator of the likelihood function in equation (1) is the value resulting from the division of equation (3) by equation (2) as shown in equation (5) below. This pseudocount value represents an approximation of the population average number of splicing junctions supporting the inclusion or exclusion of the alternative exon.

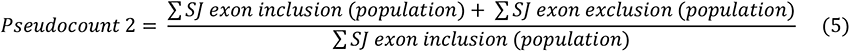

Take together, the posterior probability for a given splicing event in a given cell may be represented by the equation (6) below. A graphical representation of this model is shown in Figure 1.

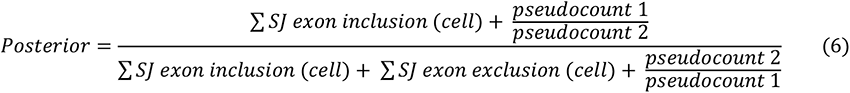

**Figure 1.**
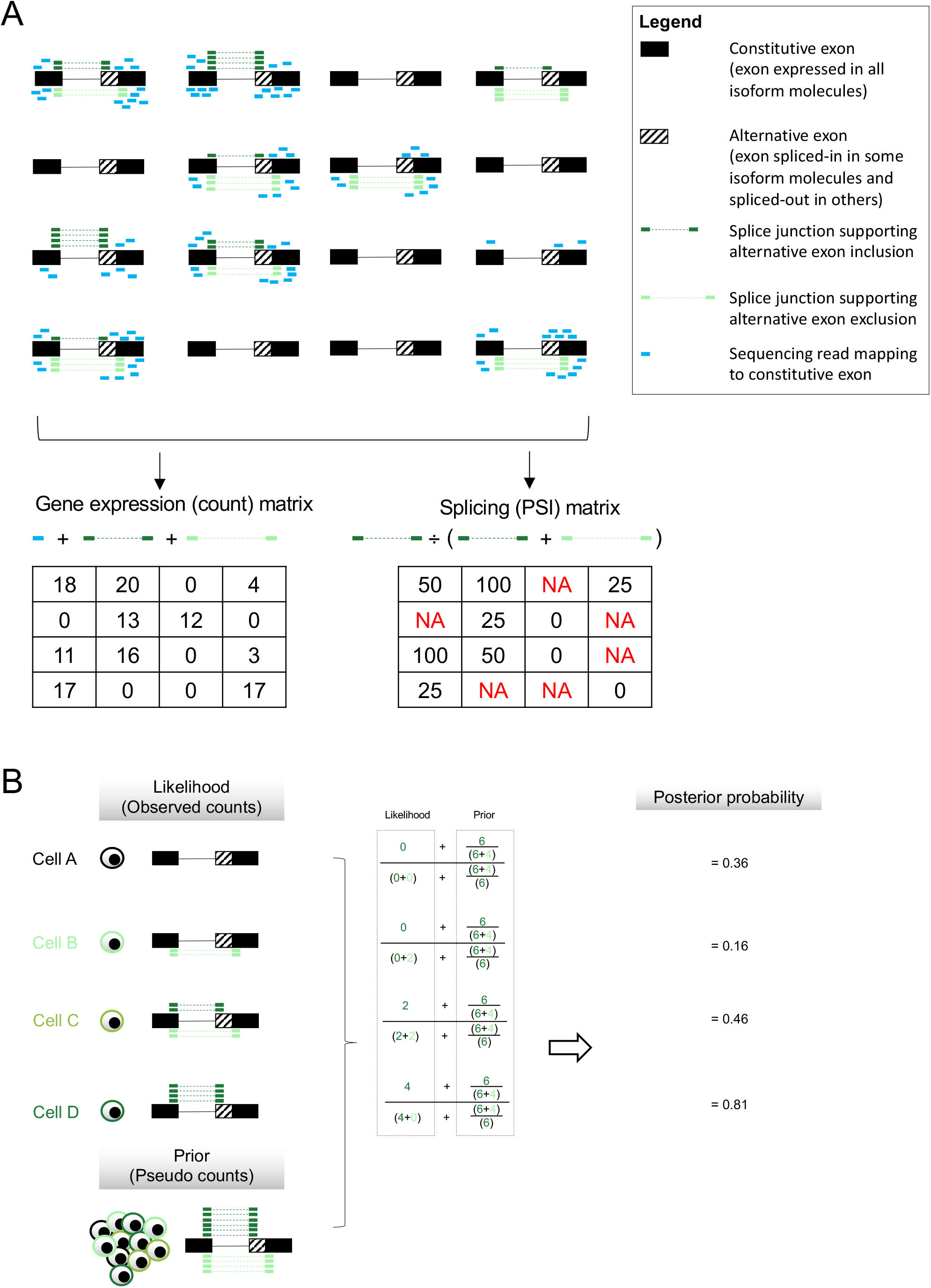
Challenges and proposed solution to dimension reduction analysis of single-cell alternative splicing dataset. **(A)** Technical dropout outs (gene expression values of 0) and low sequencing coverage prohibit robust quantification of alternative splicing (NA). Hypothetical alternative 3’ splice site across 16 cells used as an example here. The gene expression value for a given cell as indicated in the gene expression matrix is defined as the total number of reads that mapped to the constitutive exon (blue), splice junction reads that support the inclusion of the alternative exon (dark green) and splice junction reads that support the exclusion of the alternative exon (light green). On the other hand, the degree of alternative exon spliced-in as indicated in the PSI matrix is measured in PSI unit using only the observed splice junction (SJ) counts that support the inclusion (dark green) or exclusion (light green) of the alternative exon. The formula to compute PSI is Σ *SJ exon inclusion*/(Σ *SJ exon inclusion* + Σ *SJ exon exclusion*). **(B)** Combining observed splice junction counts at the single-cell level (likelihood) with observed splice junction counts derived from pseudo-bulk level (prior) yields the posterior probability. Here, the posterior probability represents the probability the exon is spliced-in. PSI: Percent spliced-in.

**Figure 2.**
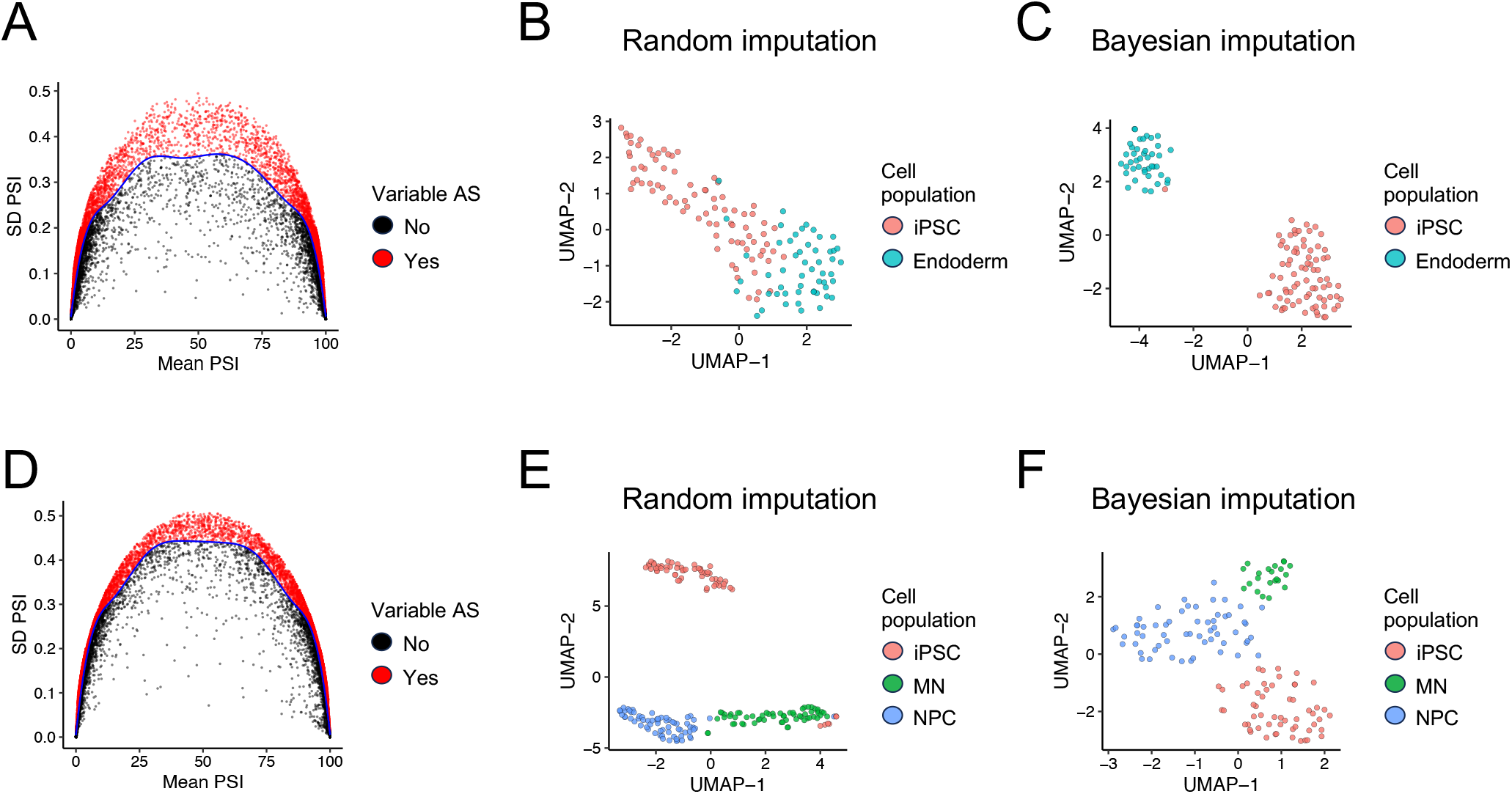
Dimension reduction analysis of transcriptomically distinct cell populations. **(A-C)** 14,630 splicing events were expressed in at least 25 cells from Linker *et al*., of which 6,358 were identified as highly variable (A) and included as features for dimension reduction analysis using the random (B) and Bayesian imputation approach (C). In total, 121 of 136 cells expressed at least 25% of splicing events specified and were included for dimension reduction analysis using the Bayesian approach in (C). **(D-F)** 16,801 splicing events were expressed in at least 25 cells from Song *et al*., of which 7,286 were identified as highly variable (D) and included as features for dimension reduction analysis using the random (E) and Bayesian imputation approach (F). In total, 136 of 190 cells expressed at least 25% of splicing events specified and were included for dimension reduction analysis using the Bayesian approach in (F). AS: Alternative splicing. iPSC: Induced pluripotent stem cells. MN: Motor neurons. NPC: Neural progenitor cells. PC: Principal component. PSI: Percent spliced-in. SD: Standard deviation.

It is noteworthy that the posterior PSI value can take any value between 0 and 1 inclusive. This is in keeping with the definition of observed PSI, i.e., the *actual rate* of alternative exon inclusion. For example, PSI of 1 indicates that the alternative exon is completely spliced-in (included) in all isoform molecules; PSI of 0 indicate that the alternative exon is completely spliced-out (excluded) in all isoform molecules; PSI of 0.5 indicate that the alternative exon is completely spliced-in in half of the isoform molecules and splice-out in half of the isoform molecules. However, unlike the observed PSI values, the posterior PSI values represent the *probability* of alternative exon inclusion.

The computation of the posterior probability is implemented in the *ComputePSI.Posterior* function of the *MARVEL* R package.

### 2.3 Data source

To benchmark and demonstrate the application of our Bayesian imputation approach for dimension reduction analysis of single-cell alternative splicing datasets, we included four datasets of increasing complexity.

The first and second datasets were derived from Linker *et al*. (Linker et al., 2019) and Song *et al*. (Song et al., 2017), respectively. Both consist of homogenous cell populations that are transcriptomically distinct from one another. Linker *et al*. consists of induced pluripotent stem cells (iPSC) and endoderm cells. Song *et al*. consists of iPSCs, motor neurons (MN) and neural progenitor cells (NPC).

The third dataset was derived from Trapnell *et al*. (Trapnell et al., 2014). This dataset consists of homogenous cell populations that are transcriptomically similar to one another. Specifically, this dataset consists of myoblast cultured and sequenced at 0-, 24-, 48-, and 72-hour timepoints.

The fourth dataset was derived from Chung *et al*. (Chung et al., 2017). This dataset consists of a mixture of cancer cells and immune cells derived from 11 breast cancer patients. Therefore, this dataset represents transcriptomically heterogenous cell populations.

### 2.4 Data processing

The pre-processed data for Linker *et al*., Song *et al*., and Trapnell *et al*. were retrieved from a recent study on single-cell alternative splicing analysis (Wen et al., 2023). From Linker *et al*. dataset, a total of 136 single cells consisting of 83 iPSCs and 53 endoderm cells were included in our analysis. From Song *et al*. dataset, a total of 190 single cells consisting of 62 iPSCs, 60 MNs, and 68 NPCs were included in our analysis. From Trapnell *et al*. dataset, a total of 327 single cell consisting of 82, 85, 88, and 72 myoblasts cultured and sequencing at 0-, 24-, 48-, and 72-hour timepoint, respectively.

From Chung *et al*. study, we downloaded the raw sequencing reads in FASTQ format from the Sequence Reads Archive (SRA; accession number SRP067248). Pre- processing and quality control were performed as previously described (Wen et al., 2023). In total 490 single cells were included in our analysis consisting of 305 tumour cells and 185 non-tumour cells. The tumour compartment consists of 70 ER+ cells, 124 HER2+, 25 ER+ HER2+ and 86 TNBC cells whereas the non-tumour compartment consists of 80 B cells, 50 T cells, 36 myeloid cells, and 19 stromal cells.

### 2.5 Dimension reduction analysis

Linear dimension reduction analysis was performed using principal component analysis (PCA) with the *PCA* function implemented by the *FactoMineR* R package. The eigenvalues for each principal component (PC) were subsequently computed using *get_eigenvalue* function implemented by the *factoextra* R package. Non-linear dimension reduction analysis was performed using Uniform Manifold Approximation and Projection (UMAP) using the *umap* function implemented by the *umap* R package. PCA, eigenvalue calculation, and UMAP may be performed using the wrapper *RunPCA* function of the *MARVEL* R package.

For each dimension reduction analysis, we first performed linear dimension reduction analysis using PCA and plotted PC1 and PC2 on the 2D space or PC1, PC2, and PC3 on the 3D space. If the cells clustered by their respective cell population, we then proceeded with non-linear dimension reduction analysis using UMAP. But if non-linear dimension reduction analysis did not segregate the different cell populations, we then presented the results from PCA.

For integrated splicing and gene expression analysis, we first performed PCA for splicing and gene features, separately. After which, we plotted the percentage of variance explained by the first 50 PCs using the *ElbowPlot* function of the *MARVEL* R package. The first PCs that explained the most variance, i.e., PC1 up the n^th^ PC that reached the plateau of the elbow plot, for the splicing and gene PCs, were combined as the final set of features for UMAP.

We used multivariate analysis of variance (MANOVA) to assess the separation of the different cell populations on the reduced dimension space (Lipovetsky, 2015). This enabled objective evaluation, on top of visualise appraisal, of the separation of the different cell populations. MANOVA was performed with the *manova* function available as part of the *stats* R package. Specifically, the cell group label was defined as the outcome in the model. Principal components 1 and 2 were defined as the independent variables when the clusters were presented with 2D PCA. Principal components 1, 2, and 3 were defined as the independent variables when the clusters were presented with 3D PCA. Finally, UMAP-1 and −2 axes were defined as the independent variables when the clusters were presented with UMAP.

### 2.7 Identification of highly variable splicing events

Highly variable splicing events were used for the initial unsupervised clustering of single cells. We first modelled the standard deviation (SD) of the PSI values as a function of the average PSI values using the P-splines approach implemented by the *gam* function of the *mgcv* R package. We then applied the model to compute the predicted SD PSI for a given mean PSI. To visualise the model, the SD PSI values (y-axis) were plotted against the mean PSI values (x-axis) for each splicing event (Liu & Zhang, 2020), and the predicted SD PSI values were represented as a smoothed curve. Splicing events with observed SD PSI greater than the predicted SD PSI were classified as highly variable and included as features for downstream analysis. We required the splicing event to be expressed (supported by at least 10 splice junction reads) in at least 25 cells for inclusion for the modelling. Both identification and plotting of the highly variable splicing events may be performed using the *IdentifyVariableEvents* function of the *MARVEL* R package.

### 2.8 Differential splicing and gene expression analysis

Differential splicing analysis was performed using the D test statistics (DTS) approach which assessed the differences in overall PSI distribution between two cell populations (Dowd, 2020).

For Trapnell *et al*. dataset, we performed differential splicing analysis for all possible pairs of cell population as previously described for homogenous cell populations (Wen et al., 2023). Only splicing events expressed in at least 25 cells in both cell groups were included for differential splicing analysis. Differentially spliced events were defined as *P* values < 0.05 and |ΔPSI| > 5%. Cells expressing at least 25% of differentially spliced events were included for dimension reduction analysis.

For Chung *et al*. dataset, we performed differential splicing analysis of each cell population against all other cell populations combined as previously described for heterogenous cell populations (G. Wang et al., 2022). Only splicing events expressed in at least 10 cells in both cell groups were included for differential splicing analysis. Differentially spliced events were defined as *P* values < 0.05 and |ΔPSI| > 10%. Cells expressing at least 10% of differentially spliced events were included for dimension reduction analysis.

Differential gene expression analysis between two cell groups was performed using the Wilcoxon rank-sum test. Only genes expressed in at least 25 and 10 cells in either cell group for Trapnell *et al*. and Chung *et al*. dataset, respectively, were included for differential gene expression analysis. *P* values were adjusted for multiple testing using Benjamini-Hochberg procedure. Differentially expressed genes were defined as FDR < 0.10 and |log2 fold change| > 0.5 were included for dimension reduction analysis.

Both differential splicing and gene expression analysis may be performed using the *CompareValues* function of the *MARVEL* R package.

### 2.9 Pathway enrichment analysis

Genes that are differentially spliced across the different cell populations in Trapnell *et al*. dataset (see section 2.8: Differential splicing and gene expression analysis) were subjected to pathway enrichment analysis to identify significantly enriched biological pathways. This analysis is performed using the *BioPathways* function available as part of the *MARVEL* R package, which is simply a wrapper function for the *clusterProfiler* R package required for the analysis. A list of significantly enriched biological processes is returned from this analysis.

For a given biological process, the pathway enrichment score is calculated. In Figure 3E for example, the “muscle filament sliding” biological process consists of 396 genes, and the number of differentially spliced genes that overlapped with this gene list is 6. The gene ratio is then calculated as 6/396. On the other hand, there were 18,866 protein-coding genes in the human genome included as background in this analysis, and the number of differentially spliced genes that overlapped with this gene list is 39. The background ratio is then calculated as 39/18,886. The pathway enrichment score is finally calculated as the ratio of gene ratio to background ratio, resulting in a score of 7.34 as indicated on the x-axis.

**Figure 3.**
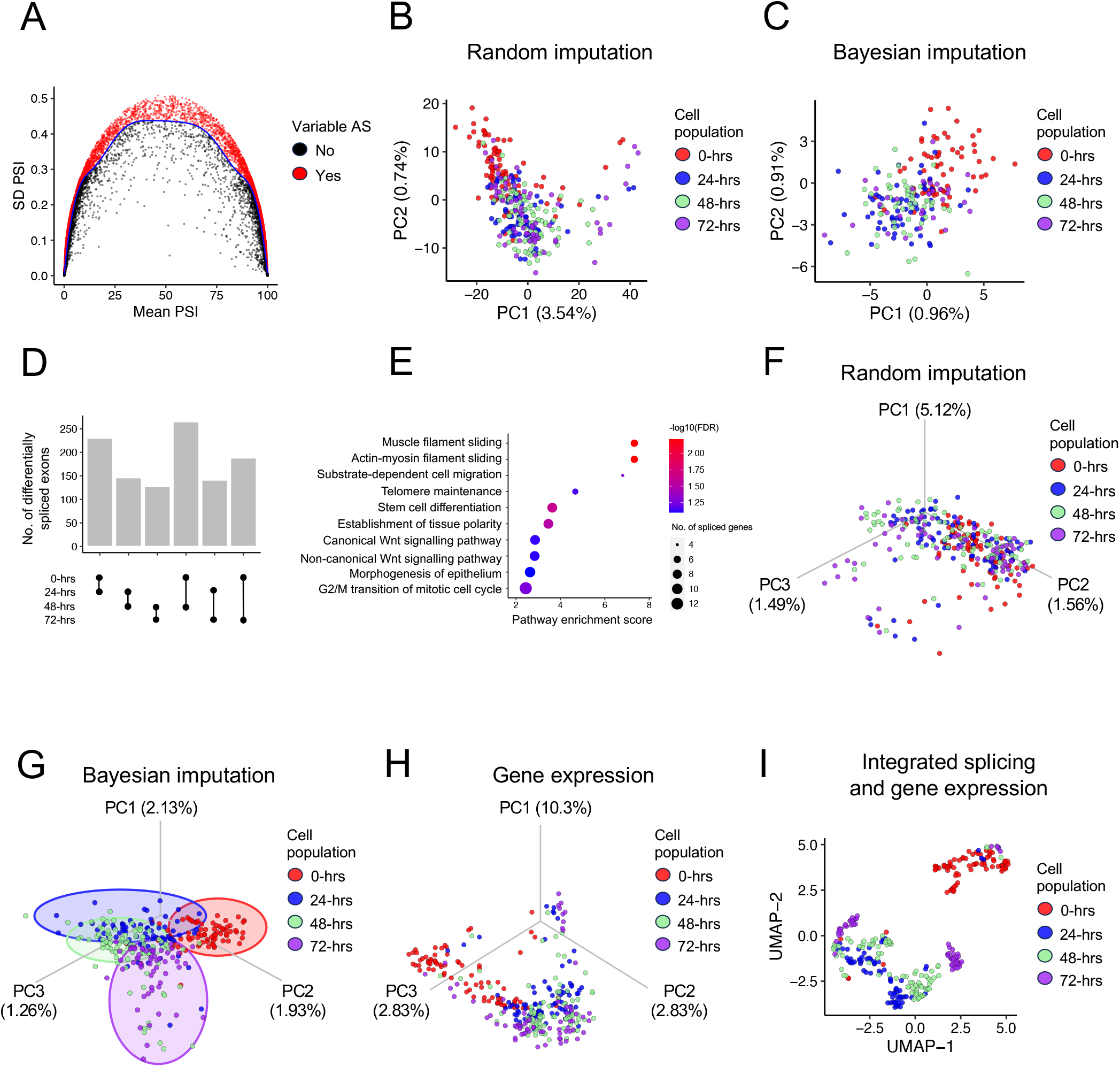
Dimension reduction analysis of transcriptomically similar cell populations. **(A-C)** 15,590 splicing events were expressed in at least 25 cells from Trapnell *et al*., of which 6,727 were identified as highly variable (A) and included as features for dimension reduction analysis using the random (B) and Bayesian imputation approach (C). In total, 212 of 327 cells expressed at least 25% of splicing events specified and were included for dimension reduction analysis using the Bayesian approach in (C). **(D-E)** Differential splicing analysis identified 603 unique splicing events (D) and belonged to genes related to muscle cell differentiation and maturation (E). **(F-G)** Differentially spliced events included as features for reduction dimension analysis using the random (F) and Bayesian imputation approach (G). In total, 295 of 327 cells expressed at least 25% of splicing events specified and were included for dimension reduction analysis using the Bayesian approach in (G). **(H)** Differentially expressed genes included as features for dimension reduction analysis. **(I)** Principal components from (G) and (H) used as features for dimension reduction analysis. AS: Alternative splicing. hrs: Hours. PC: Principal component. PSI: Percent spliced-in. SD: Standard deviation.

## 3 Results

### 3.1 Problem statement and proposed solution

In general, the PSI value is calculated as the percentage or fraction of reads supporting the inclusion (splicing in) of the alternative exon divided by the total number of reads supporting the inclusion or exclusion (splicing out) of the alternative exon. Therefore, the PSI value may be considered as probability of the alternative exon being included (spliced-in). However, computing PSI based on observed splice junction counts alone will lead to some cells whose isoforms have no or low coverage to have missing values, and consequently preclude them from downstream dimension reduction analysis (Figure 1A).

The solution proposed to overcome the issue of missing PSI values is a Bayesian approach. The PSI value computed using the observed splice junction read counts is the likelihood component of the Bayesian model. However, there is uncertainty in PSI quantification of isoforms with low or no splice junction reads. Here, we leveraged the population-level splice junction read counts as the prior in the Bayesian model. By combining both the likelihood and prior, we obtained the final PSI value as a posterior probability (Figure 1B).

Therefore, for isoforms with moderate-to-high sequencing coverage, the single-cell splice junction read counts (likelihood) will be the major contributor to the posterior probability. For isoforms with low sequencing coverage, the population-derived splice junction read counts (prior) will be the major contributor to the posterior probability. For isoforms with no sequencing coverage, the posterior probability will be totally dependent on the population-derived splice junction read counts (prior).

The end point of the Bayesian imputation is a PSI matrix, where the rows represent the splicing events and columns represent single cells, without missing values and therefore suitable for downstream dimension reduction analysis.

### 3.2 Benchmark and application on transcriptomically distinct cell populations

We hypothesised that both random and Bayesian imputation approaches will be able to readily separate homogenous cell populations that are transcriptomically distinct from one another. Indeed, unsupervised clustering of cell populations derived from Linker *et al*. based on highly variable splicing events successfully distinguished induced pluripotent stem cells (iPSC) from endoderm cells when the random or Bayesian imputation approach was used (*P*MANOVA < 2.2 x 10^-16^ for both approaches; Figures 2A-C) as previously shown (Wen et al., 2023).

We next assessed the ability of random and Bayesian imputation approaches to distinguish three cell populations. Unsupervised clustering of cell populations derived from Song *et al*. based on highly variable splicing events successfully distinguished iPSC, motor neurons (MN), and neural progenitor cells (NPC) when the random or Bayesian imputation approach was used (*P*MANOVA < 2.2 x 10^-16^ for both approaches; Figures 2D-F) as previously shown (Song et al., 2017).

In conclusion, we successfully reproduced the results of previously reported dimension reduction analysis that implemented the random imputation approach. We additionally shown, perhaps unsurprisingly, that our Bayesian imputation approach similarly distinguished these transcriptomically distinct cell populations. Therefore, Linker *et al*. and Song *et al*. datasets served as positive controls or gold standards for the optimisation of our Bayesian imputation approach.

### 3.3 Benchmark and application on transcriptomically similar cell populations

To date, dimension reduction analysis of single-cell alternative splicing has not been demonstrated beyond transcriptomically similar cell populations. Therefore, we next sought to assess the ability of the random and Bayesian imputation approaches to distinguish transcriptomically similar cell populations. To this end, we utilised Trapnell *et al*. dataset that consists of myoblast cell populations that were cultured and sequenced at four different time points (0-, 24, 48-, and 72-hour). While both approaches were able to distinguish the cell populations (*P*MANOVA < 2.2 x 10^-16^ for both approaches), the separation of clusters was largely driven by the separation of 0-hour myoblast population from myoblast of other timepoints as indicated by the attenuated *P* values when the 0-hour myoblast were excluded from analysis (random imputation: *P*MANOVA = 0.026; Bayesian imputation: *P*MANOVA = 0.18). Therefore, unsupervised clustering based on highly variable splicing events did not distinguish the four different cell populations for both random and Bayesian imputation approaches (Figures 3A- C).

Next, we extracted splicing events in a supervised manner that may distinguish the different cell populations. To this end, we performed differential splicing analysis between all possible pairs of cell populations. In total, 632 unique splicing events were identified to be differentially spliced (Figure 3D). Reassuringly, the genes that constituted these splicing events were enriched in pathways related to muscle cell differentiation and development (Figure 3E). Using these differentially spliced events, we proceeded with supervised clustering of the cell populations. While the random imputation approach was able to distinguish the different cell populations (*P*MANOVA < 2.2 x 10^-16^; Figure 3F), this was largely driven by the separation of 0-hour myoblast population from the myoblast of other timepoints as indicated by the attenuated *P* value when the 0-hour myoblast population was excluded from analysis (*P*MANOVA = 0.0012).

On the other hand, the Bayesian imputation was able to distinguish all four different cell populations (*P*MANOVA < 2.2 x 10^-16^; Figure 3G). Specifically, the separation of clusters was not driven by the separation of the 0-hour myoblast population from the myoblast of other timepoints as there was no attenuation of *P* value when the 0- hour myoblast population was excluded from the analysis (*P*MANOVA < 2.2 x 10^-16^). In particular, the distinction between the most immature cell population (0-hour) versus the most mature cell population (72-hour) was the biggest, whereas the distinction between the intermediate cell populations (24-hour versus 48-hour) was relatively smaller. Nevertheless, the Bayesian imputation approach represented a notable improvement over the random imputation approach for distinguishing these cell populations.

Dimension reduction analysis based on differential gene expression analysis similarly were able to distinguish the cell populations (*P*MANOVA < 2.2 x 10^-16^; Figure 3H). This was largely driven by the separation of 0-hour myoblast population from the myoblast of other timepoints as indicated by the attenuated *P* value when the 0-hour myoblast population was excluded from analysis (*P*MANOVA = 0.0025). Specifically, differential gene expression analysis distinguished the 0- and 72-hour cell populations but had difficulty in distinguishing the intermediate 24- and 48-hour cell populations.

We hypothesised that integrating both splicing and gene-level information will be able to provide better segregation of the different cell populations compared to either information alone. To this end, we proceeded with integrating both splicing and gene-level information by combining the principal components of both splicing and gene linear dimension reduction analysis for subsequent non-linear dimension reduction analysis. We observed separation of the different cell populations (*P*MANOVA < 2.2 x 10^-16^; Figure 3I) that was not primarily driven by the separation of 0-hour myoblast population from the myoblast of other timepoints as indicated by the highly significant *P* value after removing 0-hour myoblast population from the analysis (*P*MANOVA = 2.9 x 10^-9^). Notably, we observed better distinction between the non-0-hour myoblast cell populations, in particular, among the intermediate 24- and 48-hour cell populations compared to when either splicing or gene-level information was used in our clustering analysis (Figures 3G and H).

In conclusion, transcriptomically similar cell populations required a supervised approach be distinguished from one another. In this case, differential splicing analysis was performed to identify differentially spliced events and subsequently the events were used as features for dimension reduction analysis. Further combining splicing and gene-level information led to better separation of the different cell populations compared to either splicing or gene-level information alone.

### 3.4 Benchmark and application on transcriptomically heterogenous cell populations

We have hitherto demonstrated the application of our Bayesian imputation approach on homogenous cell populations, i.e., transcriptomically presumed homogenous cell lines. Here, we extend our application to transcriptomically heterogenous cell populations derived from multiple human donors typical of medical science studies. To this end, we utilised Chung *et al*. single-cell dataset derived from breast cancer patients consisting of tumour and non-tumour cells, both compartments in turn consisting of different molecular and phenotypic subgroups of cells, respectively. Unsupervised clustering based on highly variable splicing events separated the cells into two main populations, namely tumour and non-tumour cells with both random and Bayesian imputation approaches (*P*MANOVA < 2.2 x 10^-16^ for both approaches; Figures 4A-C). Therefore, we proceeded with the analysis of the tumour and non-tumour cells separately.

**Figure 4.**
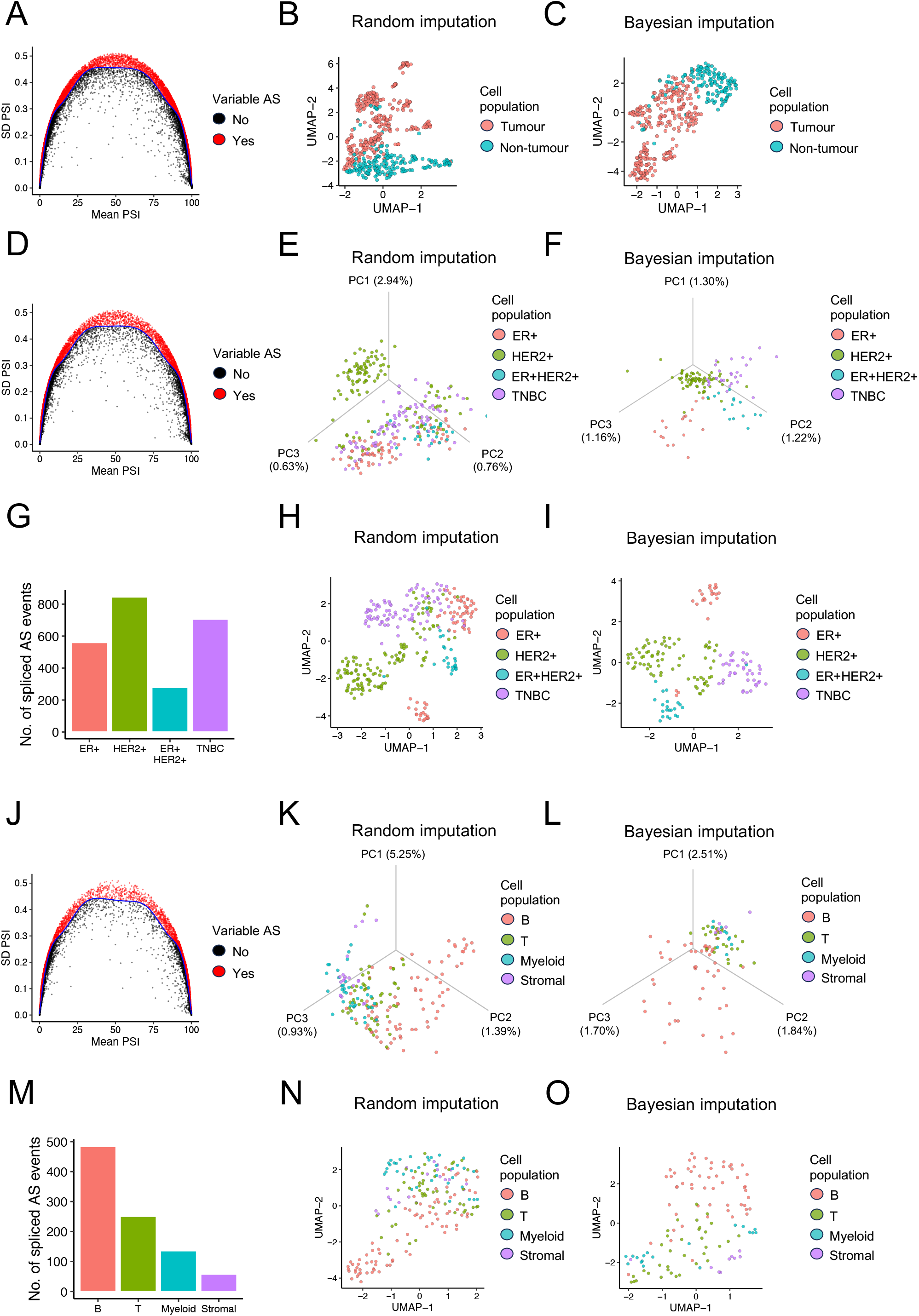
Dimension reduction analysis of transcriptomically heterogenous cell populations. **(A-C)** 23,608 splicing events were expressed in at least 25 cells from Chung *et al*., of which 11,337 were identified as highly variable (A) and included as features for dimension reduction analysis using the random (B) and Bayesian imputation approach (C). In total, 401 of 490 cells expressed at least 10% of splicing events specified and were included for dimension reduction analysis using the Bayesian approach in (C). **(D-F)** 17,986 splicing events were expressed in at least 25 tumour cells, of which 8,037 were identified as highly variable (D) and included as features for dimension reduction analysis using the random (E) and Bayesian imputation approach (F). In total, 148 of 305 cells expressed at least 25% of splicing events specified and were included for dimension reduction analysis using the Bayesian approach in (F). **(G-I)** Differential splicing analysis of tumour cells identified 1,865 unique splicing events (G) and were included as features for reduction dimension analysis using the random (H) and Bayesian imputation approach (I). In total, 159 of 305 cells expressed at least 25% of splicing events specified and were included for dimension reduction analysis using the Bayesian approach in (I). **(J-L)** 11,242 splicing events were expressed in at least 25 non-tumour cells, of which 4,527 were identified as highly variable (J) and included as features for dimension reduction analysis using the random (K) and Bayesian imputation approach (L). In total, 98 of 185 cells expressed at least 25% of splicing events specified and were included for dimension reduction analysis using the Bayesian approach in (L). **(M-O)** Differential splicing analysis of non-tumour cells identified 779 unique splicing events (M) and were included as features for reduction dimension analysis using the random (N) and Bayesian imputation approach (O). In total, 117 of 185 cells expressed at least 25% of splicing events specified and were included for dimension reduction analysis using the Bayesian approach in (O). AS: Alternative splicing. PC: Principal component. PSI: Percent spliced-in. SD: Standard deviation.

Unsupervised clustering of tumour cells based on highly variable splicing events using the random imputation approach revealed broad separation of cell populations (*P*MANOVA < 2.2 x 10^-16^; Figures 4D and E). Notably, separation of HER2+ cells from the three other cell populations (ER+, ER+HER2+, and TNBC) was observed but there were overlaps between the ER+, ER+HER2+, and TNBC cell populations. In contrast, Bayesian imputation approach revealed separation of the four cell populations, albeit with some degree of overlap between the cell populations (*P*MANOVA < 2.2 x 10^-16^; Figure 4F). Next, we investigated if supervised clustering based on differentially spliced events was able to improve the separation of the different cell populations. In total, 1,865 differentially spliced events were identified by comparing each cell population against all other cell populations (Figure 4G). Using differentially spliced events improved separation of the four cell populations compared to unsupervised clustering using highly variable splicing events for both random and Bayesian imputation approaches (*P*MANOVA < 2.2 x 10^-16^ for both approaches; Figures 4H and I).

We then proceeded with analysis of the non-tumour compartment. Unsupervised clustering based on highly variable splicing events revealed separation of B cell population from the other cell populations for both random and Bayesian imputation approaches (*P*MANOVA < 2.2 x 10^-16^ for both approaches; Figures 4J-L). We next performed the clustering in a supervised manner by using differentially spliced events as features. To this end, we identified 779 unique differentially spliced events by comparing each cell population against all other cell populations (Figure 4M). Both random and Bayesian imputation approaches identified B cells to be readily separated from the other cell populations (*P*MANOVA < 2.2 x 10^-16^ for both approaches; Figures 4N and O). Perhaps this is unsurprising given that B cells had the greatest number of differentially spliced events.

In conclusion, in the context of clustering heterogenous cell populations, both random and Bayesian imputation approaches performed comparatively well when either highly variable splicing events or differentially spliced events were used as features in the clustering analysis.

### 3.5 Benchmark PSI value estimation

BRACE employs a Bayesian approach to estimate and impute PSI values. Another single-cell splicing software that employs a Bayesian framework to estimate and impute PSI values is BRIE (Bayesian regression for isoform quantification) (Huang & Sanguinetti, 2017, 2021). We sought to benchmark and compare the PSI value estimated by these two Bayesian approaches, as well as the frequentist approach employed by MARVEL. While the Bayesian approach aims to impute missing values for splicing events with low-to-no coverage, the frequentist approach simply encodes these low coverage splicing events as missing values (Wen et al., 2023). One approach to assess PSI values is to compute the correlation of PSI values across all possible pairs of cells in a given phenotypically homogenous cell population (Hagemann-Jensen et al., 2020). To this end, we observed the cell-to-cell correlation for BRACE to be significantly higher compared to MARVEL’s frequentist approach and all three modes of estimating PSI values by BRIE (Figure 5). It is noteworthy that BRIE only estimates PSI values for SE splicing event type whereas BRACE additionally estimates PSI values for MXE, A5SS, A3SS, AFE, and ALE splicing event types. Furthermore, BRACE outperformed MARVEL’s frequentist approach for estimating PSI values for A5SS, A3SS, AFE, and ALE splicing event types. Taken together, BRACE represents a novel approach for estimating PSI values with better performance compared to published approaches.

**Figure 5.**
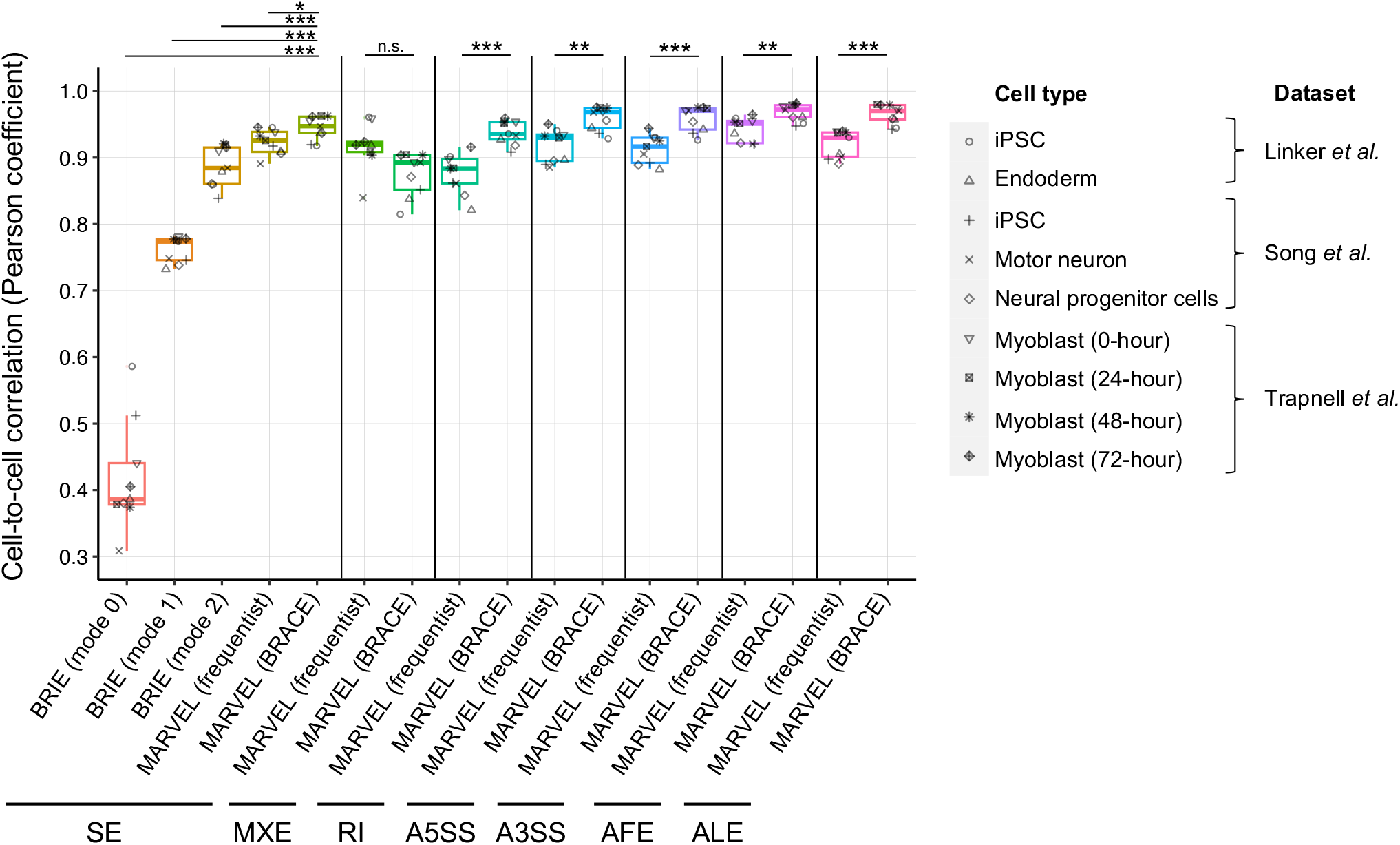
PSI value estimation of BRACE compared to published single-cell splicing software. Median cell-to-cell Pearson correlation coefficient across all pairs of cells evaluated in 9 phenotypically homogenous cell populations for PSI values computed with BRACE, BRIE, and MARVEL’s frequentist approach. Note that BRIE only computes PSI values for SE splicing events, while MARVEL’s BRACE and frequentist approaches additionally computes PSI values for MXE, A5SS, A3SS, AFE, and ALE splicing event types. For SE splicing events, only the overlapping SE events detected by both BRIE and MARVEL were included for analysis. *P* values were derived from paired t-test. A5SS: Alternative 5’ splice site. A3SS: Alternative 3’ splice site. AFE: Alternative first exon. ALE: Alternative last exon. BRIE: Bayesian regression for isoform estimation. MXE: Mutually exclusive exons. PSI: Percent spliced-in SE: Skipped-exon. *P* *** < 0.001 ** < 0.01 * < 0.05.

## 4 Discussion

In this study, we developed and demonstrated the application of a novel Bayesian-based method for dimension reduction analysis of single-cell alternative splicing datasets. To date, majority of single-cell analysis has focused on gene-level expression analysis. Nevertheless, alternative splicing represents an underappreciated and an additional layer of heterogeneity underlying gene expression profile (Wen et al., 2020). Moreover, alternative splicing may underpin biological processes in both health and disease states. Examples of the latter include diseases with aberrant splicing mechanism such as cancer (Shiozawa et al., 2018).

Differential gene expression analyses represent the cornerstone of RNA-sequencing, and dimension reduction analysis is often used for *in silico* validation of differentially expressed gene (Wen & Leong, 2019). Differentially expressed genes that are able to distinguish the different sample groups are regarded as putative true positives and lend confidence for selection of these candidate genes for further experimental characterisation and functional validation. While dimension reduction analysis of gene expression is well established, there remains significant challenges for dimension reduction analysis of alternative splicing events, especially at the single-cell level.

These challenges unsurprisingly arise from the high drop-out rate of single-cell experiments. These dropouts may be attributed to a combination technical or biological reason. The former is due to PCR amplification bottleneck whereby lowly expressed genes were not sufficiently represented in the sequencing library while the latter is due to heterogeneity within a presumed homogenous cell population whereby only a subset of cells expressed a given gene (Buen Abad Najar, Yosef, & Lareau, 2020). The result of these dropouts is reflected in the missing values prevalent in PSI matrices. While this does not hamper other aspects of single-cell alternative splicing analysis such as splicing detection, quantification, and differential splicing analysis, this issue does hamper dimension reduction analysis.

Current solution to impute these missing values include imputation of any values between 0 to 100 randomly. This solution was first proposed by Song *et al*. in the *Expedition* software (Song et al., 2017), and then further implemented by Wen *et al*. in the *MARVEL* R package (Wen et al., 2023). In both studies, the authors demonstrated that the random imputation method readily distinguished the different cell populations. While their analysis demonstrated the first proof-of-principle of dimension reduction analysis of single-cell alternative splicing datasets, it suffered several short-comings. Firstly, random imputation was not based on biology. Secondly, random imputation only included moderately-to-highly expressed splicing events, i.e., splicing events in cells supported by at least 10 sequencing reads, and thereby preclude lowly expressed, but potentially biologically informative splicing events for dimension reduction analysis. Finally, the random imputation method was demonstrated on transcriptomically distinct cell populations but its application has hitherto not been demonstrated on transcriptomically more complex cell populations.

To address these short-comings, we developed a novel Bayesian imputation approach that leveraged on population-level splicing information as a prior and combined this with single-cell splicing information regardless of the latter’s sequencing coverage. We subsequently benchmarked and applied our Bayesian method across datasets with increasing complexity, namely transcriptomically distinct, similar, and heterogenous cell populations. We were able to recapitulate the separation of transcriptomically distinct cell populations as shown in previous studies (Song et al., 2017; Wen et al., 2023). While both random and Bayesian imputation demonstrated similar performance in distinguishing heterogenous cell populations, we demonstrated our Bayesian imputation approach, but not random imputation approach, was able to distinguish transcriptomically similar cell populations. Therefore, we demonstration the applicability of our Bayesian approach across a variety of cellular context, and may represent an improvement over existing method for dimension reduction analysis of single-cell alternative splicing dataset.

The uses of dimension reduction analysis in the context of gene expression analysis include (1) identification of batch effect which enables batch correction of expression or principal component values prior to downstream analysis, (2) cell type annotation typically with Louvain method, and (3) assessing if a given list of genes (high variable or differentially expressed) is able to segregate the different cell populations (Satija et al., 2015; G. Wang et al., 2022). The focus of our study is on the third use of dimension reduction analysis, specifically, to assess if a given set of splicing events (high variable or differentially spliced) is able to segregate the different cell populations. Therefore, we anticipate our approach to be used to assess differentially spliced events identified from differential splicing analysis. In particular, the ability of these differentially spliced events to segregate the different cell populations will lend confidence to the validity of these differentially spliced events as putative true positives. Furthermore, dimension reduction analysis using differentially spliced events will aid in prioritisation of principal components that capture the most variance in the data, thus making it easier to identify the most important spliced genes that explain the differences in splicing patterns. These candidate splicing events may subsequently be prioritised for further downstream experimental studies. Therefore, it is conceivable that our approach will aid in guiding splicing-based biomarker assessment, such as cell type- or disease-specific biomarkers, in single-cell RNA-sequencing analysis.

Lastly, also notable is that our Bayesian imputation approach represents a more precise approach to estimate PSI values as demonstrated by the higher cell-to-cell correlation in PSI values compared to published Bayesian and frequentist approaches employed by BRIE and MARVEL, respectively (Huang & Sanguinetti, 2017, 2021; Wen et al., 2023). Therefore, it is conceivable that the PSI values estimated by our Bayesian approach may be applicable beyond dimension reduction analysis such as splicing pattern (modality) analysis and differentially splicing analysis.

## Data availability

All data used in this study is publicly available. The source codes for Bayesian imputation of PSI values and subsequent dimension reduction analysis are available in the MARVEL R package (version 2.0.5 onwards) on GitHub (https://github.com/wenweixiong/MARVEL).

## Acknowledgment

The author would like to thank Gonzalo Benegas for algorithm implementation, and Supat Thongjuea and Adam Mead for helpful discussions.

